# TNscope: Accurate Detection of Somatic Mutations with Haplotype-based Variant Candidate Detection and Machine Learning Filtering

**DOI:** 10.1101/250647

**Authors:** Donald Freed, Renke Pan, Rafael Aldana

## Abstract

Detection of somatic mutations in tumor samples is important in the clinic, where treatment decisions are increasingly based upon molecular diagnostics. However, accurate detection of these mutations is difficult, due in part to intra-tumor heterogeneity, contamination of the tumor sample with normal tissue and pervasive structural variation. Here, we describe Sentieon TNscope, a haplotype-based somatic variant caller with increased accuracy relative to existing methods. An early engineering version of TNscope was used in our submission to the most recent ICGC-DREAM Somatic Mutation calling challenge. In that challenge, TNscope is the leader in accuracy for SNVs, indels and SVs. To further improve variant calling accuracy, we combined the improvements in the variant caller with machine learning. We benchmarked TNscope using *in-silico* mixtures of well-characterized Genome in a Bottle (GIAB) samples. TNscope displays higher accuracy than the other benchmarked tools and the accuracy is substantially improved by the machine learning model.

## Introduction

Cancer is known to be a genomic disease, where the accumulation of genetic mutations in somatic cells results in tumorigenesis and metastasis. Genomic and other molecular analyses of tumor samples are increasingly applied to aid in clinical diagnosis and management of cancer^1,2^. Recently developed applications of next-generation sequence data include discovery of neoantigens for targeted immunotherapy and liquid biopsy for monitoring of tumor remission^3–8^. Accurate characterization of tumor genomes is essential for many of these applications, and is an active area of development.

Historically, the lack of reliable truth sets has hindered the development of accurate somatic variant callers. However, the recent release of well-characterized Genome in a Bottle (GIAB) reference samples from the National Institute of Standards and Technology (NIST) allow for improved benchmarking^9,10^. The truth sets for these samples cover approximately 80% of the human genome and are constructed using multiple technologies to minimize biases towards a specific technology. However, this process is not perfect and the truth sets may be subtly biased^11^. The GIAB truth sets generally exclude the most difficult portions of the genome, including some sites of known pathogenic mutations, leading to an overestimate of variant calling accuracy^12^. This effect is exacerbated across functional variation as purifying selection results in underrepresentation of these variants in the truth set and a relative enrichment in variant caller output. Despite these limitations, the GIAB samples remain one of the best benchmarks for assessing variant calling accuracy.

Although the GIAB datasets were originally developed for benchmarking germline variant callers, germline data from GIAB reference samples can be mixed *in-silico* to create synthetic tumor samples with true-positive somatic variants comprised of variants that are unique in one sample and absent from the other. These *in-silico* mixtures are not perfect, as some library preparation and sequencing artifacts introduced into the designated normal sample will likely be present in both the tumor and normal mixtures, the overall effect of these error modes will be underestimated. However, recent studies suggest that in practice, the magnitude of these errors is small and these *in-silico* mixtures work nearly as well as *in-vitro* mixtures for benchmarking^11^.

New tools promise to provide improved characterization of somatic variants in paired tumor-normal sample. Sentieon provides accelerated, deterministic tools for alignment, post-alignment processing and variant calling that provide matching results to BWA, MuTect and MuTect2 as well as improved algorithms, such as Sentieon TNscope^13,14^. An early engineering version of Sentieon TNscope was used in the ICGC-TCGA DREAM Mutation Calling 6 challenge and is at the top of the performance leaderboard across snvs, indels, and structural variants^15^.

By default, TNscope uses rule-based variant filters for identifying false-positive variant calls, similar to MuTect2 and most other somatic variant callers. In practice, these rule-based filters do not fully utilize the information in the variant annotations provided by the variant callers. For improved variant filtration, we trained a random forest model using a small subset of variant calls from a mixture of two different GIAB samples. Application of the trained model to the full variant callsets produced by TNscope results in a substantially improved accuracy across a range of depths and variant allele frequencies.

In this study, we benchmarked TNscope with and without the machine learning model for variant filtration, using *in-silico* mixtures of well-characterized GIAB samples. For comparison, we also benchmarked Sentieon TNsnv and Sentieon TNhaplotyer. TNsnv and TNhaplotyper provide matching results to MuTect and MuTect2 respectively, but without downsampling for improved accuracy and consistency. Our results suggest that Sentieon TNscope has improved accuracy over both TNsnv and TNhaplotyper for somatic variation across a range of sample depths and allele fractions, and that the variant calling accuracies are further improved by the machine learning filtering.

## Sentieon TNscope

TNscope is a haplotype-based variant caller that follows the general principles of the mathematical models first implemented in the GATK HaplotypeCaller and MuTect2^16–18^. This includes active region detection, assembly of haplotypes from the reference and local read data using a de Bruijn-like graph, pair-HMM for calculation of read-haplotype likelihoods followed by genotype assignment. Similar to MuTect2, TNscope evaluates haplotypes jointly in the tumor and normal samples when normal samples are available, significantly increasing precision for somatic variant detection.

A number of improvements have been made in TNscope’s mathematical model to increase recall and precision for somatic variation. The computational efficiency of the Sentieon tools allows for lower thresholds for triggering active regions, facilitating a more complete evaluation of sites of potential variation. Further, the detected active regions are typically of higher quality as TNscope uses statistics to trigger active regions rather than hard cutoffs. Local assembly is improved, resulting in more frequent identification of the correct variant haplotype. Genotyping is improved with a novel quality score and modified nonparametric statistical tests for filtering false-positive variant candidates. TNscope also includes a number of novel variant annotations that can be used for improved variant filtration. Tumor-only mode is supported with a panel of normal samples. TNscope’s computational efficiency and no-downsampling make it an ideal haplotype-based variant caller for detection of low-level somatic events in high-depth targeted sequencing data.

## Methods

### Generation of *in-silico* mixtures, post-alignment processing, and variant calling

*In-silico* mixtures for somatic variant benchmarking were generated from 300x Illumina HiSeq 2500 data sequenced at NIST. Each sample in a tumor-normal pair was generated with data from separate flowcells, avoiding the possibility of using data repeatedly in both the tumor and normal samples. Within the designated flowcells for the tumor and normal samples, a subset of fastq files were selected that would best approximate the desired target coverage for the *in-silico* mixture (Supplementary Table 1 and Supplementary Table 2).

Each selected fastq pair was aligned independently to the human reference genome b37 (hs37d5) using Sentieon enhanced BWA-MEM. Sentieon enhanced BWA-MEM has about 2X speedup to BWA-mem while providing identical results. Read group information including library id, and read group ID (consisting of library id, flowcell id and lane number) were added to each BAM during alignment on the basis of information derived from the NIST FTP site. The read group sample name was added as either the sample name or a concatenation of the tumor and normal sample names depending upon whether the fastq were being used to compose the normal or tumor samples, respectively. Post-alignment processing steps including duplicate marking, indel realignment, and BQSR were performed with standard settings (Supplementary Appendix 1).

For TNsnv, TNhaplotyper and TNscope, variant calling was performed with the option “--min_init_tumor_lod 1.0” to lower the emission threshold for variants. This option does not change the set of PASS variants, but it is useful when generating the precision recall plots. For the TNscope runs used with the machine learning model (referred to as TNscope-model for the remainder of the text), a set of parameters were tuned to lower the thresholds for active region detection and variant emission, allowing the model to obtain higher sensitivity than would be possible with default settings. Additionally, with TNscope-model, post-variant calling filtering was applied using the trained machine learning model while other FILTER tags were removed (Supplementary Appendix 1).

### Variant evaluation

Variant evaluation used version 3.3.2 of the GIAB truth sets for samples HG001, HG002, HG004 and HG005. For a pair of samples designated as a tumor or normal sample, variants present in the tumor sample and absent from the normal sample were identified with bcftools isec, part of the SAMtools package^19^. High confidence regions for the sample pair were intersected and overlapping intervals were merged using BEDtools^20^. While attention has been paid to different haplotype representation when comparing a callset to a truth set, little attention has been paid to consistent haplotype representation across truth sets. Within the truth sets we find evidence of differential haplotype representation that is not accounted for by bcftools. To remove potential errors due to differential haplotype representation, we force-call variants that are supposedly unique to the tumor sample in the normal sample. Variants that are successfully called with an alternate allele depth of greater than nine or an alternate allele fraction of greater than 0.1 are removed from consideration through subtraction of the locus from the high-confidence region using BEDtools subtract. Please see Supplementary Appendix 1 for more details.

Evaluation of variant caller accuracy was performed with RTGtools vcfeval (version 3.7.1) separately for SNVs and indels^21^. For the generation of accuracy benchmark tables, default settings were used with the addition of the “--squash-ploidy” parameter.

To generate the precision-recall plots, additional settings were used. For TNsnv, the output VCF was annotated with the reported TLOD from the call stats file. If the call stats file indicated that “fstar_tumor_lod” was the only applicable filter, then the FILTER field in the VCF would be changed from REJECT to PASS. For TNhaplotyper and TNscope, the “t_lod_fstar” FILTER was removed from the FILTER field using bcftools. For TNsnv, TNhaplotyper and TNscope, comparison with the truth set was performed using RTGtools vcfeval with the additional settings “--squash-ploidy --vcf-score-field=INFO.TLOD”. For TNscope-model, an annotation was constructed from the TLOD annotation output by the variant caller and the ML_PROB annotation output by the machine learning model. In cases where ML_PROB equals 1.0, ROC_VAL is set equal to 10 plus the TLOD, otherwise ROC_VAL was set equal to ML_PROB. For comparison with the truth set using RTGtools vcfeval, the additional settings were “--squashploidy --vcf-score-field=INFO.ROC_VAL --all-records”.

## Results

The variant calling process necessarily involves statistically modeling next-generation sequence data. While the major signals of genetic variation are readily identified with straightforward statistical models, the high dimensionality of these data make application of standard statistical techniques for variant filtration sub-optimal. To address this issue, we sought to improve accuracy using supervised machine learning methods. First, we created a synthetic paired tumor-normal sample using an *in-silico* mixture of HG001 (normal) and HG002 (tumor) at various tumor/normal depths and tumor purity fractions (Supplementary Table 3). TNscope was used to call variants on these samples with sensitive settings (see Methods). These sensitive settings lower thresholds for active region detection and variant emission and include a local *de novo* assembly process that is more robust in repetitive regions. Variants in the resulting callsets were then classified as true-positive or false-positive using the GIAB truth set. We then trained a random forest classifier for both SNPs and indels on this output using the hyperparameters described in Supplementary Table 4 on a random subset of the classified variants. A major strength of the random forest model is the high accuracy that may be obtained from relatively small training datasets. Our model was trained on only 100,000 variants from each mixture (approximately 2.5% of all variants present in the high-confidence regions of the training samples). Evaluation of model performance on the held-out data revealed high accuracy across these samples (Supplementary Table 5) and suggests that some of the novel annotations in TNscope are partially correlated with empirical variant quality.

For a benchmark dataset, we created an *in-silico* mixture by adding reads from HG005 (tumor) into HG004 (normal) at various allele fractions and depths. Tumor samples were paired with a corresponding pure normal sample (Supplementary Table 6). Variants were called from these *insilico* mixtures with TNsnv, TNhaplotyper, TNscope with default settings, and TNscope with sensitive setting and variant filtration using the machine learning model (Figure 1, Supplementary Figure 1, and Supplementary Table 7). With default settings, TNscope displayed consistently higher sensitivity and F1-score than TNhaplotyper and TNsnv. The trained machine learning model substantially improved F1-score beyond TNscope with default settings, especially for indels, likely due to the model’s ability to capture interaction between annotations and perform more effective parameter tuning.

**Figure 1.**
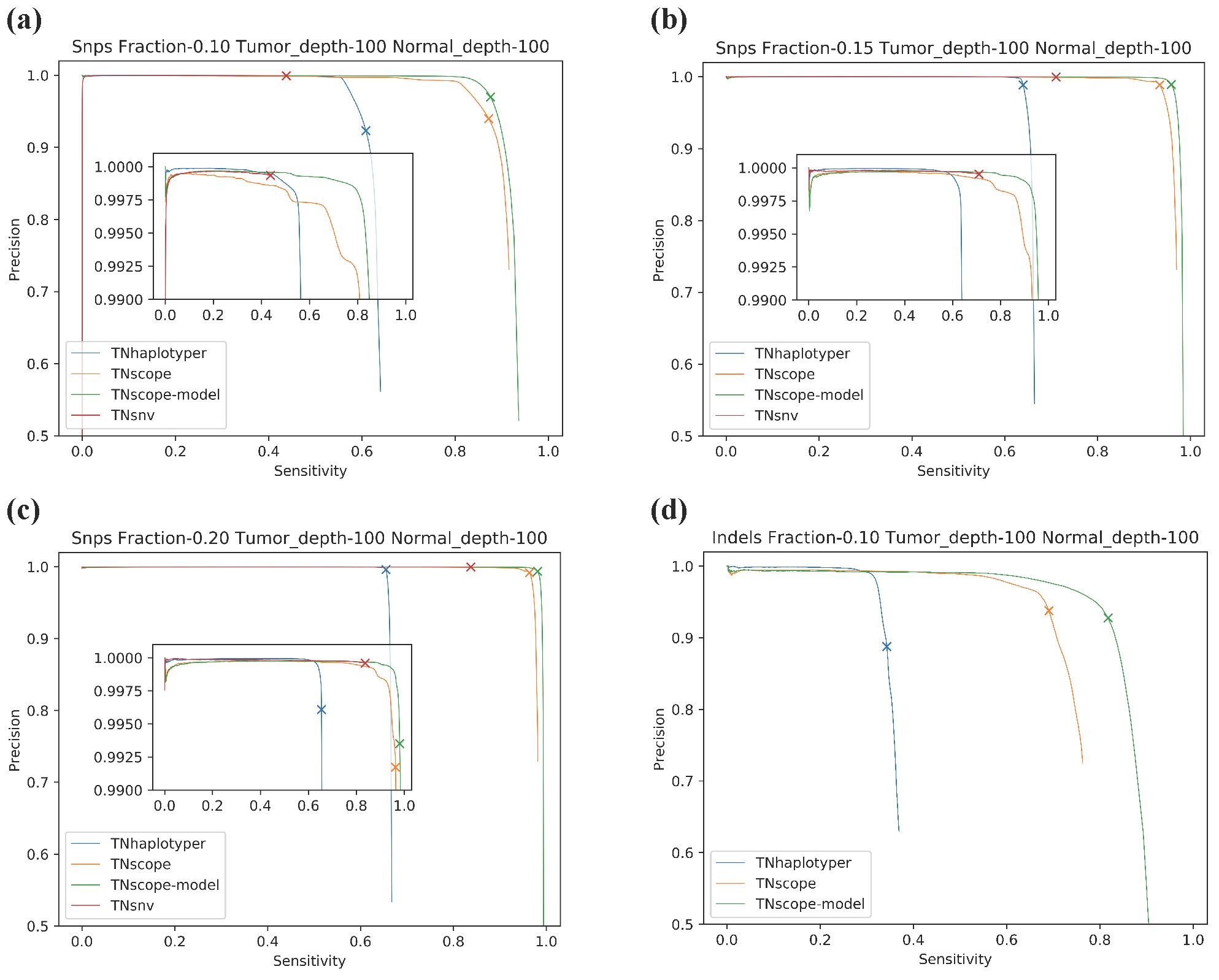
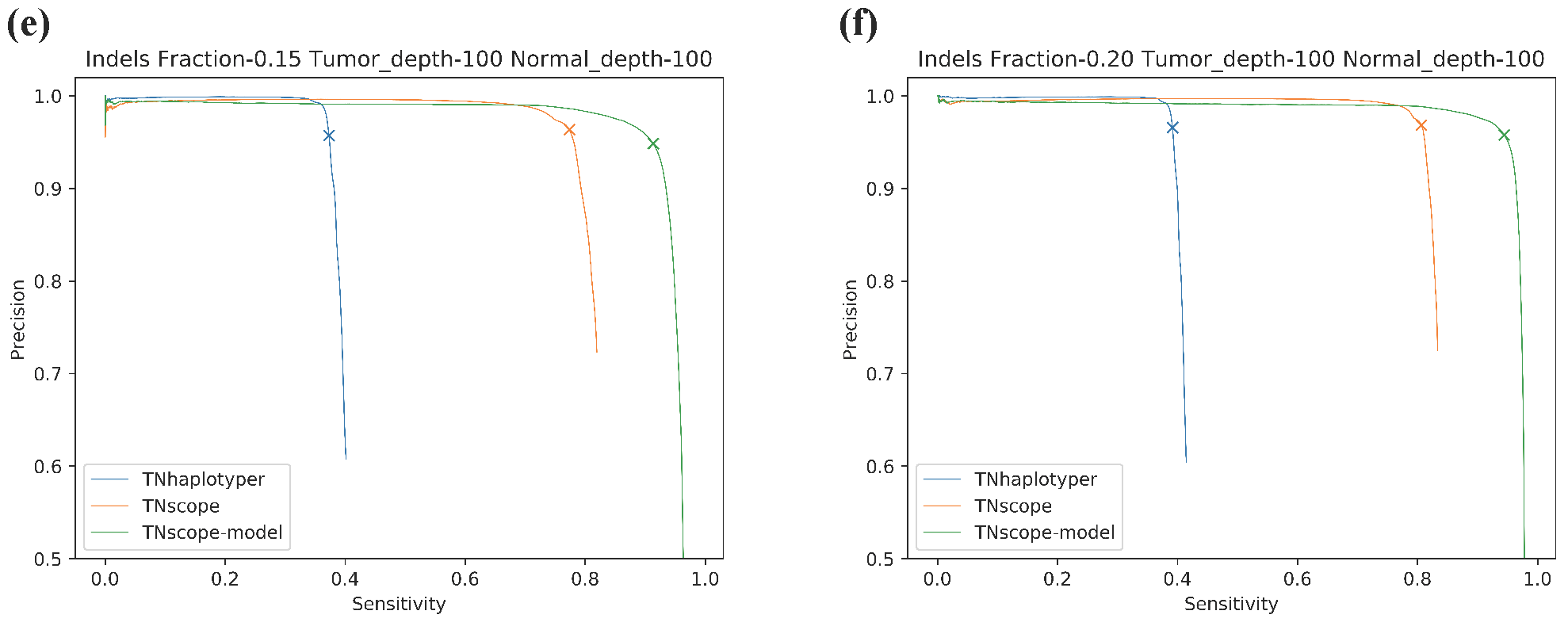
Precision-recall curves for TNsnv, TNhaplotyper, and TNscope with two settings. “TNscope” is the TNscope variant caller with default settings while “TNscope-model” is TNscope with sensitive settings followed by machine learning for variant filtration. All samples are at 100x depth in the tumor and normal samples with tumor sample purities from 0.10 to 0.20. “X” marks the maximum F1-score for each algorithm. (a-c) SNVs. (d-f) indels.

## Discussion

In this manuscript, we present Sentieon TNscope and benchmark TNscope against industry-standard somatic variant callers. TNscope has the highest accuracy across the benchmarked tools and variant filtration using our trained random forest classifier further improves variant calling accuracy. Given the high accuracy of TNscope with a small number of reads supporting the variant allele and its ability to process large datasets without downsampling, we believe that TNscope will be especially useful for detection of low-level somatic variation. The machine learning model also provides a single ensemble annotation that can effectively be used for filtering, eliminating the need to manually tune variant annotation filters.

The use of haplotype-based variant calling for variant candidate detection and machine learning for variant filtration combines the strengths of both approaches. Computationally efficient statistical models can be used for variant candidate detection, improving feature engineering and greatly reducing feature space, while machine learning filtration can be used to provide improved accuracy.

## Acknowledgements

We thank Justin Zook and the NIST team for developing the truth sets. We thank Jonathan Pevsner and Sean Cho of Johns Hopkins University for helpful discussions regarding benchmarking experiments.

## Author Contributions

D.F., and R.A. designed, performed and evaluated the experiments. R.P. developed, implemented and tested the machine learning model. D.F. wrote the manuscript with input from all authors.

## Data Availability

The publically available data used in these experiments can be downloaded from the Genome in a Bottle FTP site (ftp://ftp-trace.ncbi.nlm.nih.gov/giab/ftp/)

## Supplementary Figures

**Supplementary Figure 1.**
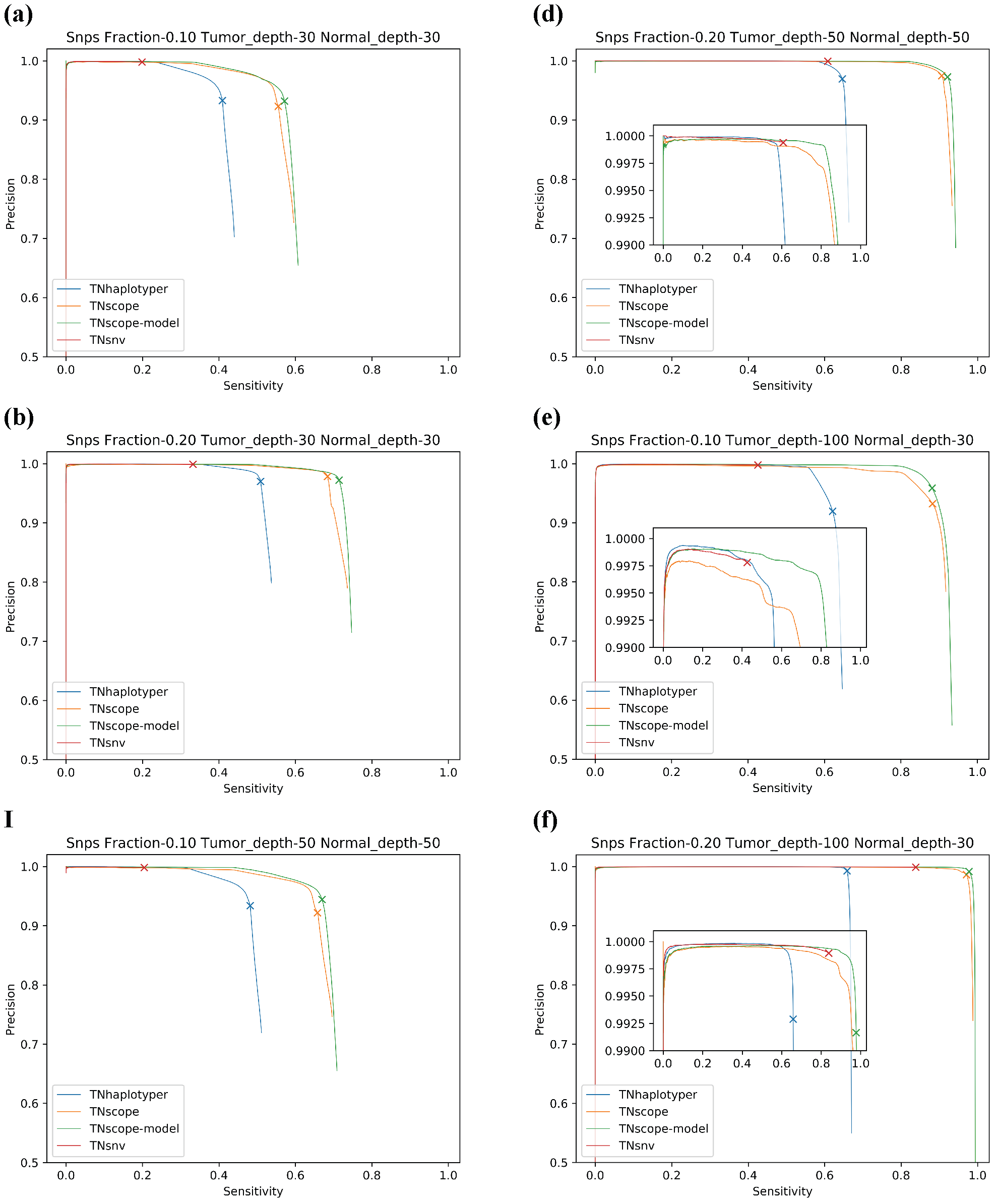
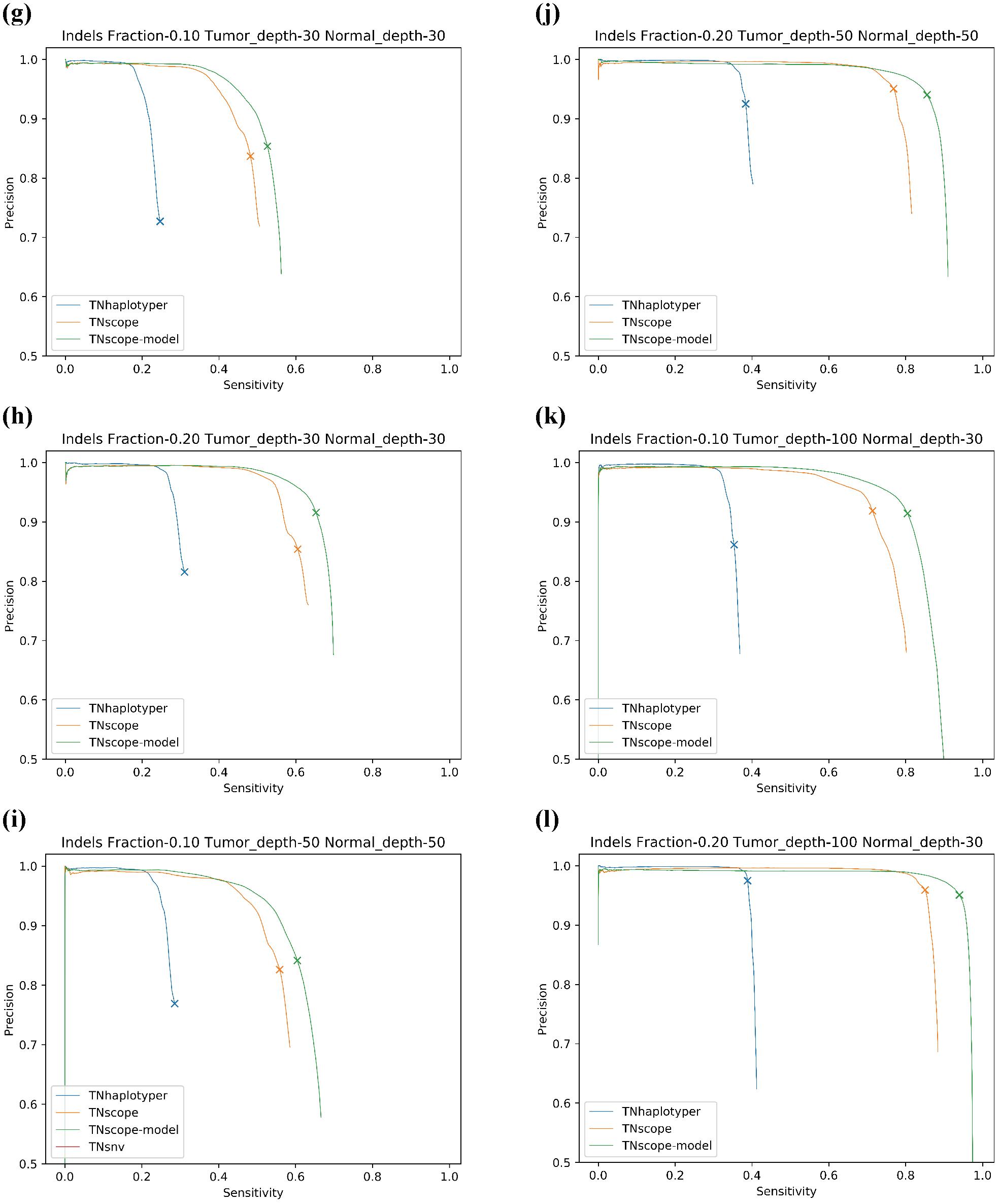
Precision-recall curves for TNsnv, TNhaplotyper, and TNscope with two settings.

“TNscope” is the TNscope variant caller with default settings while “TNscope-model” is TNscope with sensitive settings followed by machine learning for variant filtration. “X” marks the maximum F1-score for each algorithm. (a-f) SNVs. (g-l) indels.

## Supplementary Tables

**Supplementary Table 1- Fastq files used to create *in-silico* mixtures for training the machine learning model.**

**Supplementary Table 2- Fastq files used to create *in-silico* mixtures for the benchmark dataset.**

**Supplementary Table 3- Sample depths and tumor purities used during model training.** Values in cells are denote the tumor sample purity in the mixture fractions. The normal sample is HG001 with the tumor sample HG002.

**Supplementary Table 4- Hyperparameters used during model training**. “max_depth”: The maximum depth of leaves in a tree. “min_sample_count”: the minimum number of variants required to split a leaf node. “use_surrogates”: enables surrogates for missing data. “max_categories”: cluster some features to find a suboptimal split (important for limiting training runtime). “cal_var_importance”: Enable calculation of variant importance. “nactive_vars”: The number of randomly selected features to evaluate at the tree. “forest_accuracy”: Stop training early if accuracy is close to perfect. “Training_Percentage”: Fraction of the supplied data to use for model training.

**Supplementary Table 5- RTGtools vcfeval results on held-out data.** Includes both SNPs and indels.

**Supplementary Table 6- Sample depths and tumor purities used in the benchmark sample.** Values in cells are denote the tumor sample purity in the mixture fractions. The normal sample is HG004 with the tumor sample HG005.

**Supplementary Table 7- RTGtools vcfeval results on the benchmark sample.** The evaluation results for each of the *in-silico* mixtures used in the benchmark.

## Supplementary Files

**Supplementary Appendix 1- Example commands.**

